# Voltage-Gated Potassium Channel Kv1.3 as a Therapeutic Target for Pancreatic Ductal Adenocarcinoma

**DOI:** 10.1101/2021.02.03.429505

**Authors:** Weiwei Li, Gregory C. Wilson, Magdalena Bachmann, Jiang Wang, Andrea Mattarei, Cristina Paradisi, Michael J. Edwards, Ildiko Szabo, Erich Gulbins, Syed A. Ahmad, Sameer H. Patel

## Abstract

The mitochondrial voltage-gated potassium channel, Kv1.3, has been emerged as an attractive oncologic target but its function in pancreas cancer (PDAC) is unknown. In this study we evaluated tissue expression of Kv1.3 in resected PDAC from 55 patients and tumor inhibition in orthotopic mouse models using the recently developed Kv1.3 inhibitors *PCARBTP* and *PAPTP*. Immunohistochemistry of 55 human PDAC specimens showed that all tumors expressed Kv1.3 with 60% of tumor specimens having high Kv1.3 expression. In pancreas tumor models (Pan02 cells injected into C57BL/6 mice), *PCARBTP* and *PAPTP* treatment resulted in tumor reductions of 87% and 70%, respectively. When combined with gemcitabine/abraxane, this increased to 95% and 80% without resultant organ toxicity. *In vivo* models indicated *PCARBTP*-mediated cell death occurred through the p38-MAPK pathway. In vitro-generated resistant clones to *PCARBTP* escaped cell death through upregulation of the anti-oxidant system as determined using SWATH-MS analysis. These data show Kv1.3 is highly expressed in resected human PDAC and the use of novel mitochondrial Kv1.3 inhibitors combined with cytotoxic chemotherapies might be novel, effective treatment for PDAC.

## Introduction

Despite the utilization of more aggressive systemic chemotherapy regimens, pancreas ductal adenocarcinoma (PDAC) remains a devastating disease and is the third leading cause of cancer related mortality. ^1^ In 2020, pancreas cancer will affect approximately 57,600 patients in the US and the incidence is expected to rise.^2^ Surgery remains the only option for cure but unfortunately, only 15-20% of patients are candidates for resection and five year overall survival remains less than 20% with surgery alone. Apart from surgery and chemotherapy, few effective treatment options exist. Immunotherapy, which has shown dramatic results in many gastrointestinal and cutaneous malignancies, thus far has shown minimal benefit in PDAC. One of the barriers to treatment is the tumor microenvironment, which is rich in immunosuppressive cells.^3^ These regulatory T cells, myeloid derived suppressor cells, and tumor associated macrophages (TAMs) allow tumor cells to evade identification and are involved in the development of resistance.^3, 4^ Novel treatment strategies are desperately needed.

Voltage-dependent K^+^ channels (Kv) are a superfamily of ubiquitously expressed membrane proteins that are involved in maintaining membrane resting/action potentials, cell proliferation, immune activation, and cell death.^5^ Kv1.3 is a specific voltage-dependent K^+^ channel located mainly in the plasma and inner mitochondrial membranes (mitoKv1.3). First discovered in the plasma membrane of human T lymphocytes, Kv1.3 is also found in tumor and immune cells where they regulate proliferation as well as apoptosis and are aberrantly expressed in malignancies.^6–8^ Kv1.3 is inhibited by many chemically unrelated compounds such as small-molecule organic compounds, and venom-isolated oligopeptides.^9–15^ Our group has recently developed two specific mitoKv1.3 inhibitors that prevalently and specifically target the mitochondrial channel by virtue of a positively charged triphenylphosphonium group. These inhibitors (*PCARBTP* and *PAPTP*) were shown to selectively kill cancer cells but not normal healthy cells through a reactive oxygen species (ROS)-mediated cell death involving the respiratory chain complex I. ^16–18^

*In vitro* and *in vivo* treatment of metastatic human pancreas cancer cell lines with mitochondrial Kv1.3 inhibitors resulted in cell death and reduced tumor growth but not complete eradication.^16^ Furthermore, the clinical impact of Kv1.3 in resected human PDAC specimens is not known. Therefore, in this study we sought to evaluate tissue expression of Kv1.3 in resected PDAC, and assess tumor growth inhibition using novel inhibitors of mitoKv1.3 in conjunction with cytotoxic chemotherapy.

## Results

### Kv1.3 is Highly Expressed in Resected Human Specimens and PDAC cell lines

Immunohistochemical (IHC) evaluation was performed on tissues from 55 patients who were diagnosed at a relatively early stage, in time to undergo resection for PDAC (Figure 1A). Median patient age was 68 years old with 42% of patients being female and 86% being white followed by 9% black. A whipple operation (pancreaticoduodenectomy) was performed in 69.1% of patients, distal pancreatectomy in 29.1%, and total pancreatectomy in 1.8%. On final pathology, 29.1% of specimens had poorly differentiated tumors, 90.9% had perineural invasion, and 56.4% lymphovascular invasion. Positive lymph nodes were found in 83.6% of patients. American Joint Committee on Cancer (AJCC 8^th^ edition) staging showed 52.7% were stage 2, 34.5% stage 3, and 12.7% were stage 1.

**Figure 1.**
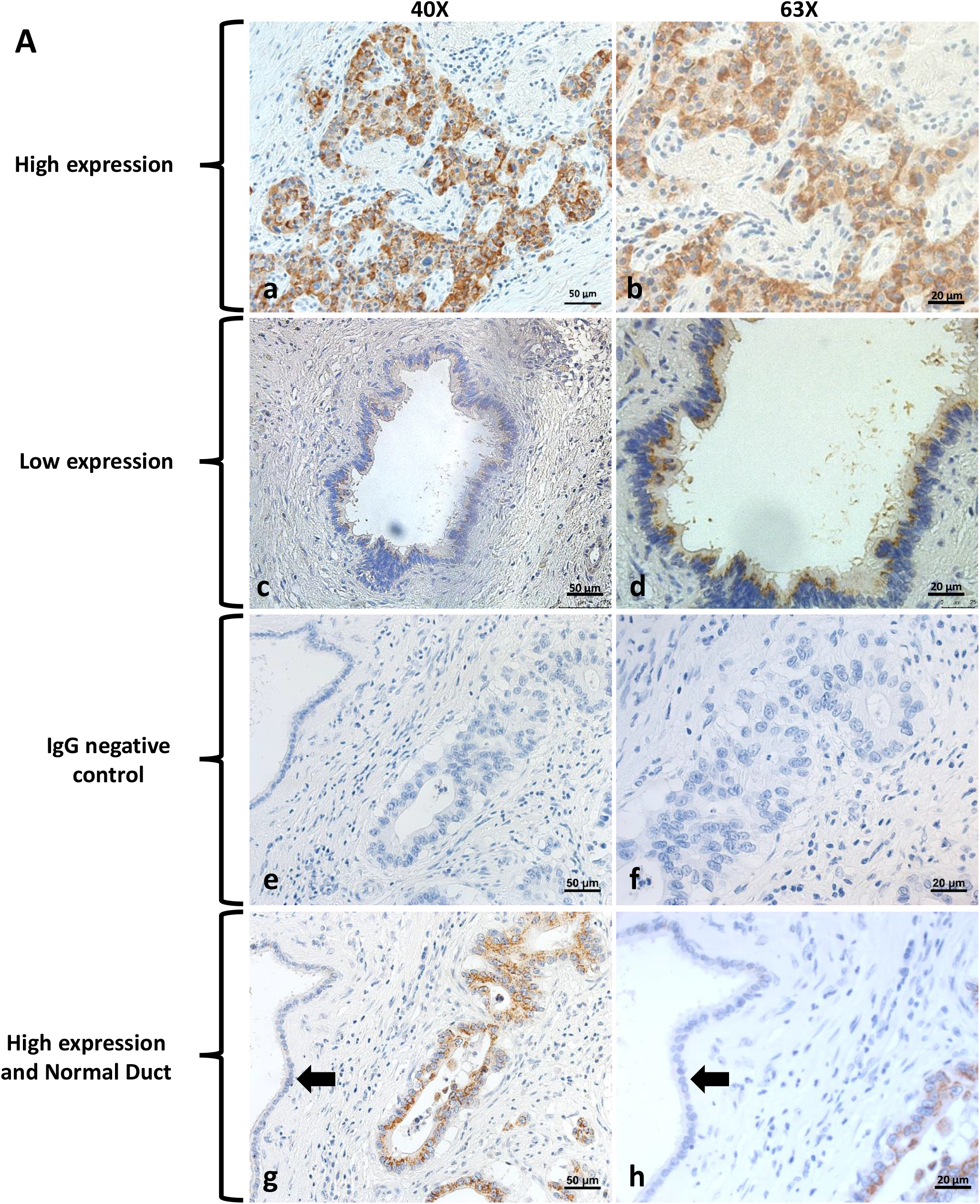
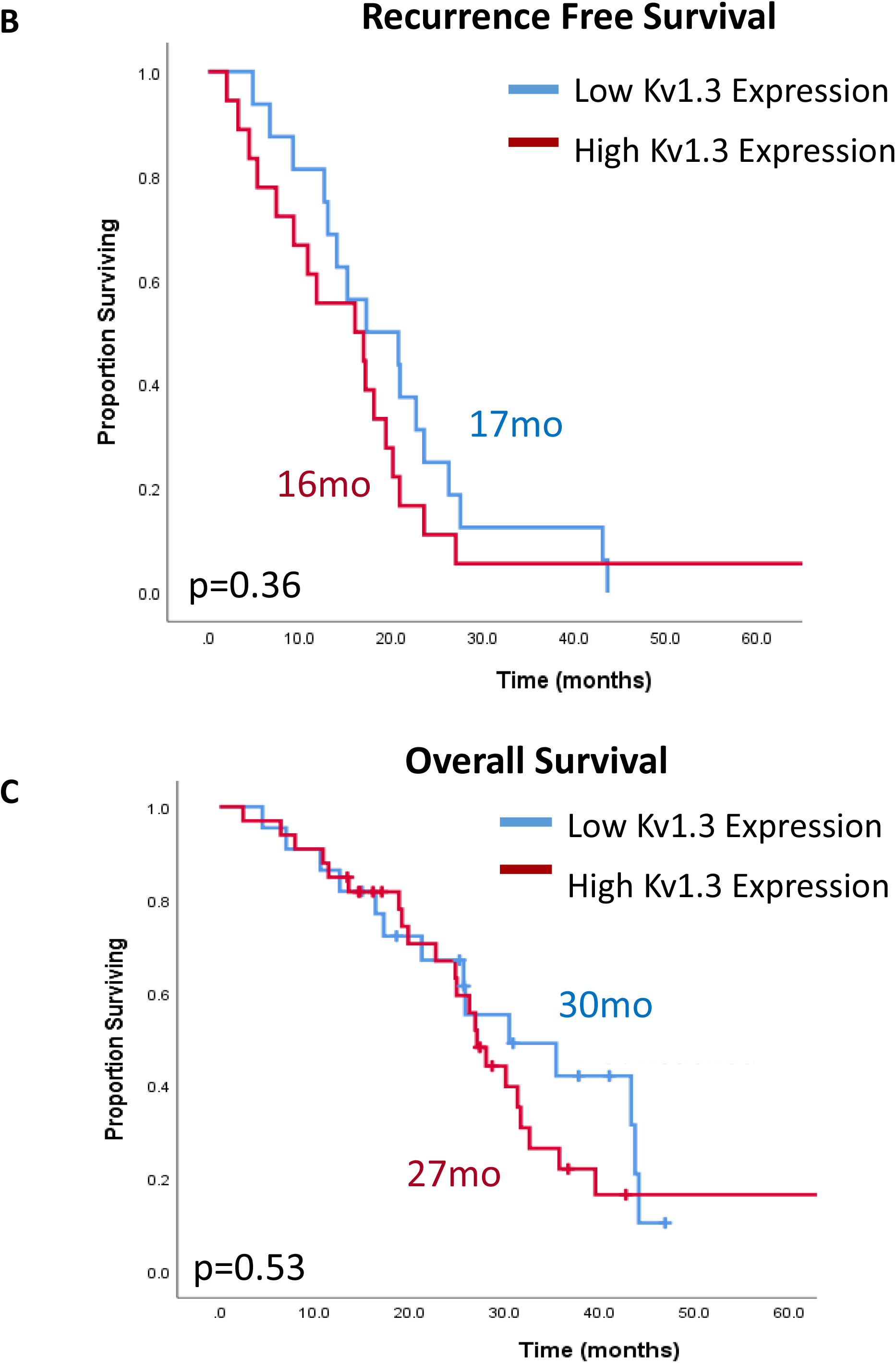
Expression of Kv1.3 in human pancreatic cancer tissue and pancreatic cell lines. A. Expression of Kv1.3 in human pancreatic tissue. Panel a, high expression level of Kv1.3 in human pancreatic cancer tissue. (n=33, scale bar=50 μm). Panel b, magnification of panel a, scale bar=20μm. Panel c, low expression level of Kv1.3 in human pancreatic cancer tissue. (n=22, scale bar=50 μm). Panel d, magnification of panel c, scale bar=20μm. Panel e, IgG control with magnification (Panel f). Panel g and h show high expression of Kv1.3 and an example of a normal pancreas duct (black arrow). B, Recurrence free survival and C, overall survival based on Kv1.3 expression in resected human pancreatic cancer tissues.

Kv1.3 IHC staining showed that 96.4% (n=53) of specimens exhibited expression. Based on percentage and intensity of stain, we found that 60.0% of tumor specimens had high expression. Only 8.3% of normal pancreas specimens had expression of Kv1.3. Over a median follow up of 28.0 months, 67.9% of patients developed a recurrence and 65.5% of patients died. When stratified by Kv1.3 expression, median recurrence free survival was 16 months with high Kv1.3 expression versus 17 months with low expression (p=0.36) (Figure 1B). Median overall survival was 27 months with high Kv1.3 expression and not reached 30 months for low Kv1.3 expression (p=0.53) (Figure 1C). High expression of Kv1.3 in tumor specimens was not associated with having a positive lymph node, poor differentiation, perineural invasion, or lymphovascular invasion (p>0.05).

### *PCARBTP* and *PAPTP* Reduced Pancreatic Ductal Adenocarcinoma Tumor Size in an Orthotopic Mouse Model

Since no studies have been performed on PDAC in immune-competent mice with mitoKv1.3 inhibitors, tumor growth was examined here in the setting of treatment with specific mitoKv1.3 inhibitors in an orthotopic model injecting mouse Pan02 cells. Six days post tumor injection, 1003D mini-pumps (MICRO-OSMOTIC PUMP MODEL 1003D, Reservoir Volume, 100 μl, 1.0 μl per hour, 3 days) were implanted filled with 100 μL of 50% DMSO, 25 mM *PCARBTP* and 50 mM *PCARBTP* respectively at a flow rate of 1 or 2 nmol/h/gbw (Table 1). 3 days after implantation, mini-pumps were replaced once fulfilled with same concentration drug at D10, tissues were collected at D14. The tumor volume and mass treated with lower dose of *PCARBTP* were reduced by 60% (n=7) and 50% (n=7), the tumor volume and mass treated with the higher dose *PCARBTP* were reduced by 72% (n=7) and 68% (n=7), respectively (Figure 2A). There were no significant changes in body weights of the mice with drug treatment from the pre-orthotopic injection body weights (p=0.84 for 1 nmol/h/gbw, p=0.22 for 2 nmol/h/gbw, Figure 2B). Representative tumor images are shown in Figure 2C.

**Table 1.** Treatment Regimen of Figure 2 A-C.

| Day 0 | Day 7-12 | Day 14 |
| --- | --- | --- |
| Before tumor injection | 1003D mini-pump duration | Tumor collected |

**Figure 2.**
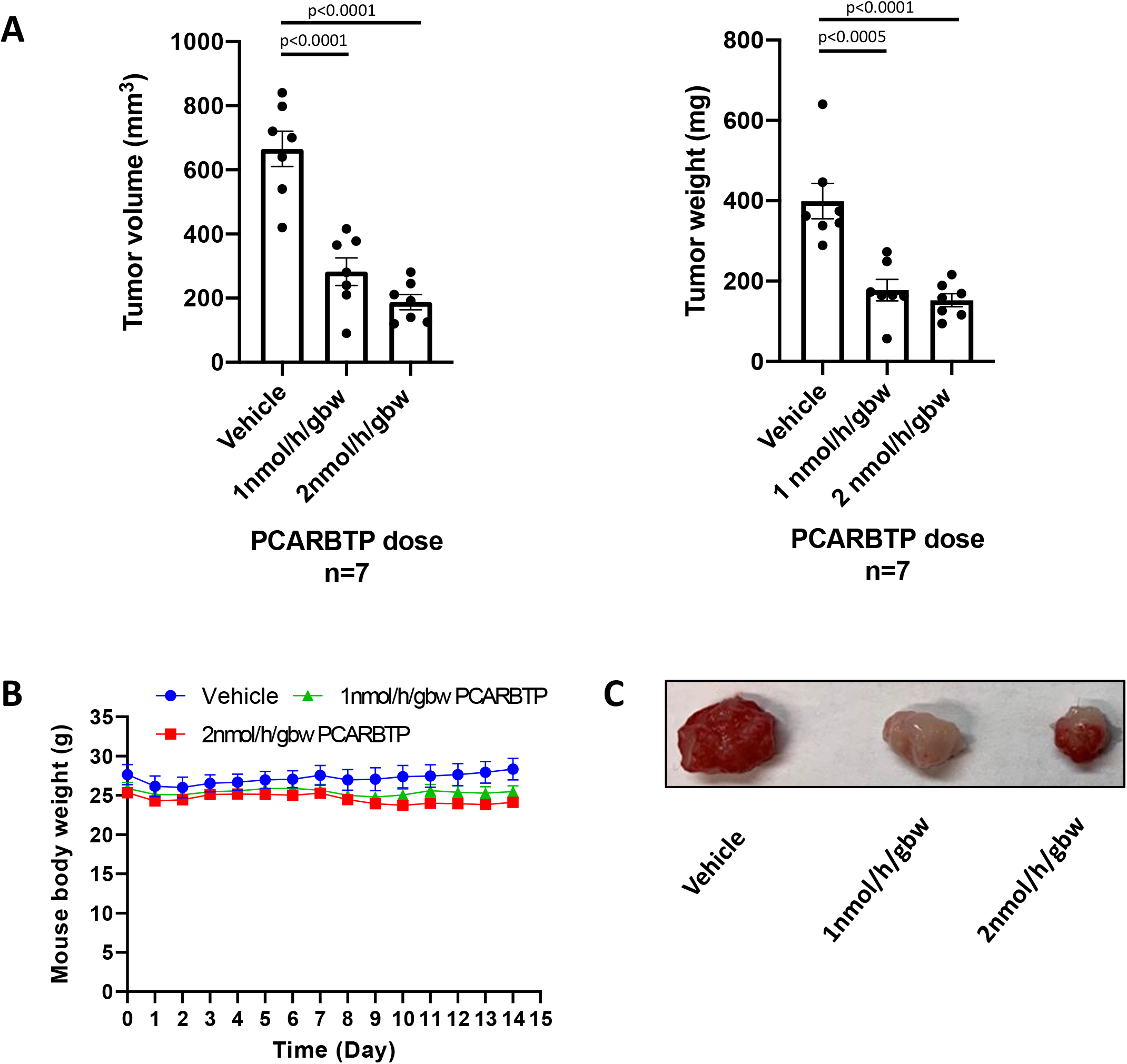
*PCARBTP* reduced pancreatic tumor size in mouse models through mini-pump administration. A. *PCARBTP* reduced mouse pancreatic tumor volume orthotopically significantly by mini-pump application method, dose of *PCARBTP* was 1nmol/h/gbw and 2nmol/h/gbw. *PCARBTP* reduced mouse pancreatic tumor weight orthotopically significantly by mini-pump application method, n=7. B. Bodyweights change of the mice during *PCARBTP* treatment were no difference (p=0.84 for 1nmol/h/gbw and p=0.22 for 2nmol/h/gbw). C. Representative pictures of the mouse orthotopically injected tumors, n=7.

An additional drug application method was used with intraperitoneal injection of *PCARBTP* and P*APTP* beginning at 6 days post the tumor injection, and tumors were collected at day 12 post tumor injection. 15 nmol/g body weight and 20 nmol/g body weight of *PCARBTP* were injected every other day for 3 doses (Table 2). Likewise, 5 nmol/g body weight *PAPTP* was injected every other day for 3 doses (Table 3). The tumor volume and mass treated with 15 nmol/gbw *PCARBTP* were reduced by 87% and 87% (n=8). The tumor volume and mass treated with 20 nmol/gbw *PCARBTP* were reduced by 88% and 90% (n=8) (Figure 3A). Although there were decreases in body weight from the pre-orthotopic weight after *PCARBTP* treatment at both the 15 and 20 nmol/gbw doses (p=0.02 and p=0.006, respectively), no mice had greater than a 20% decrease body weight (Figure 3B). Representative images are shown in Figure 3C. The tumor volume and mass treated with 5 nmol/gbw *PAPTP* were reduced by 70% (n=8) and 64% (n=8), respectively (Figure 4A). There were no significant changes in body weights of the mice with *PAPTP* treatment from the pre-orthotopic injection body weights as shown in Figure 4B (p=0.17). Representative tumors are shown in Figure 4C.

**Table 2.**
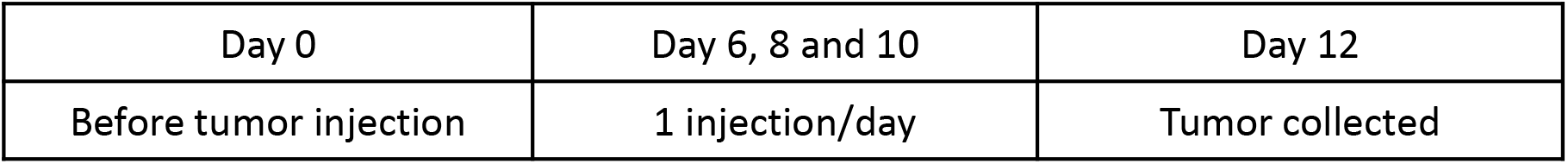
Treatment Regimen of Figure 3 A-C.

|  |  |  |
| --- | --- | --- |
| Day 0 | Day 6, 8 and 10 | Day 12 |
| Before tumor injection | 1 injection/day | Tumor collected |

**Table 3.**
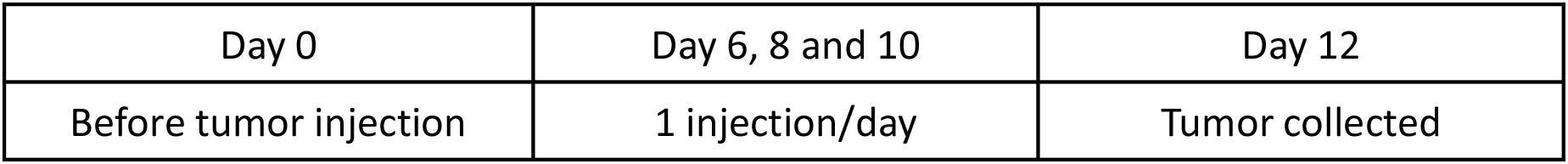
Treatment Regimen of Figure 4 A-C.

| Day 0 | Day 6, 8 and 10 | Day 12 |
| --- | --- | --- |
| Before tumor injection | 1 injection/day | Tumor collected |

**Figure 3.**
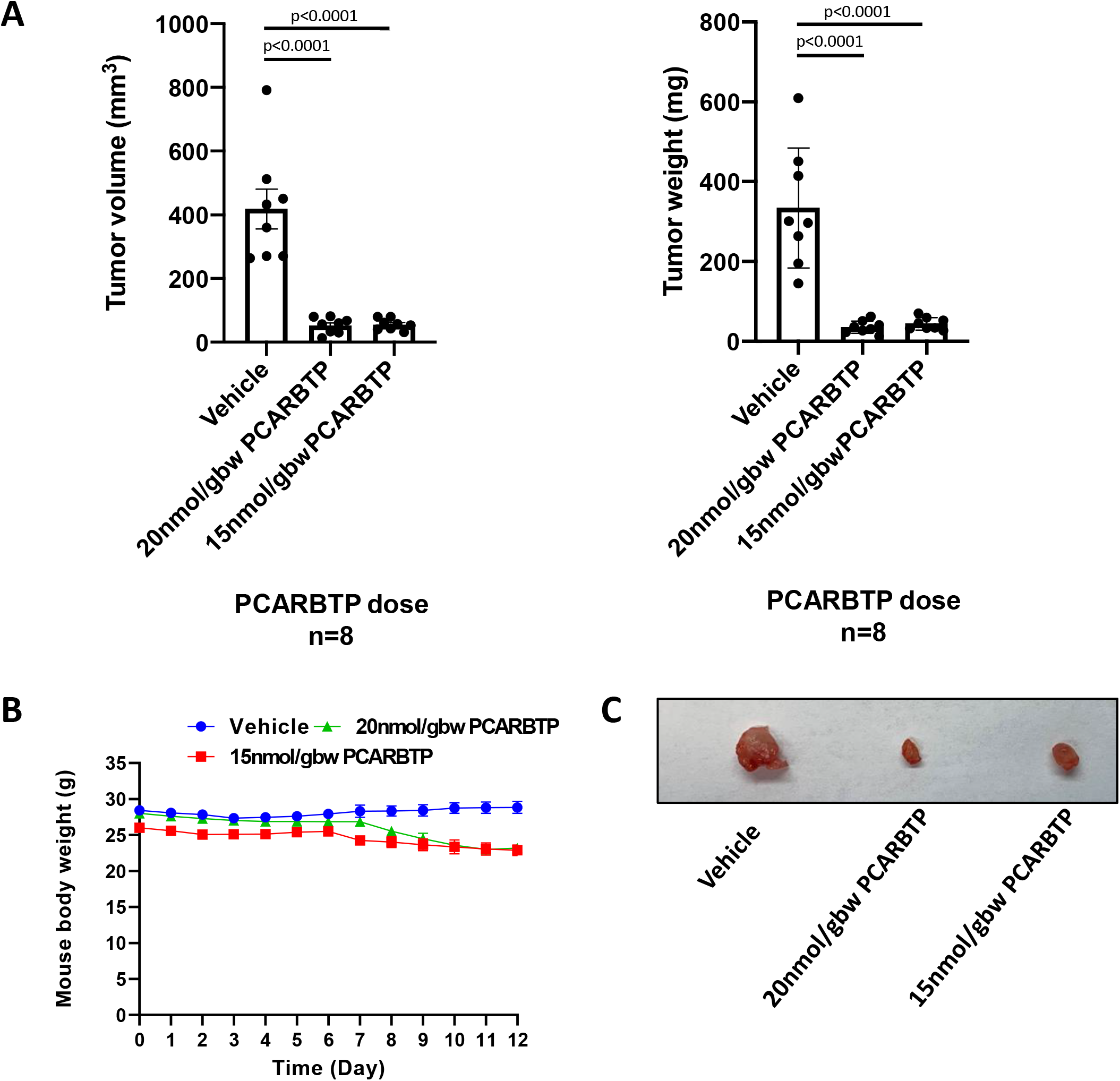
*PCARBTP* reduced pancreatic tumor size in mouse models through intraperitoneal administration. A. *PCARBTP* reduced mouse pancreatic tumor volume orthotopically significantly by intraperitoneal administration, n=8. *PCARBTP* reduced mouse pancreatic tumor weight orthotopically significantly by intraperitoneal administration. B. Bodyweights change of the mice during *PCARBTP* treatment were less than 20% (p<0.05). C. Representative pictures of the mouse orthotopically injected tumors.

**Figure 4.**
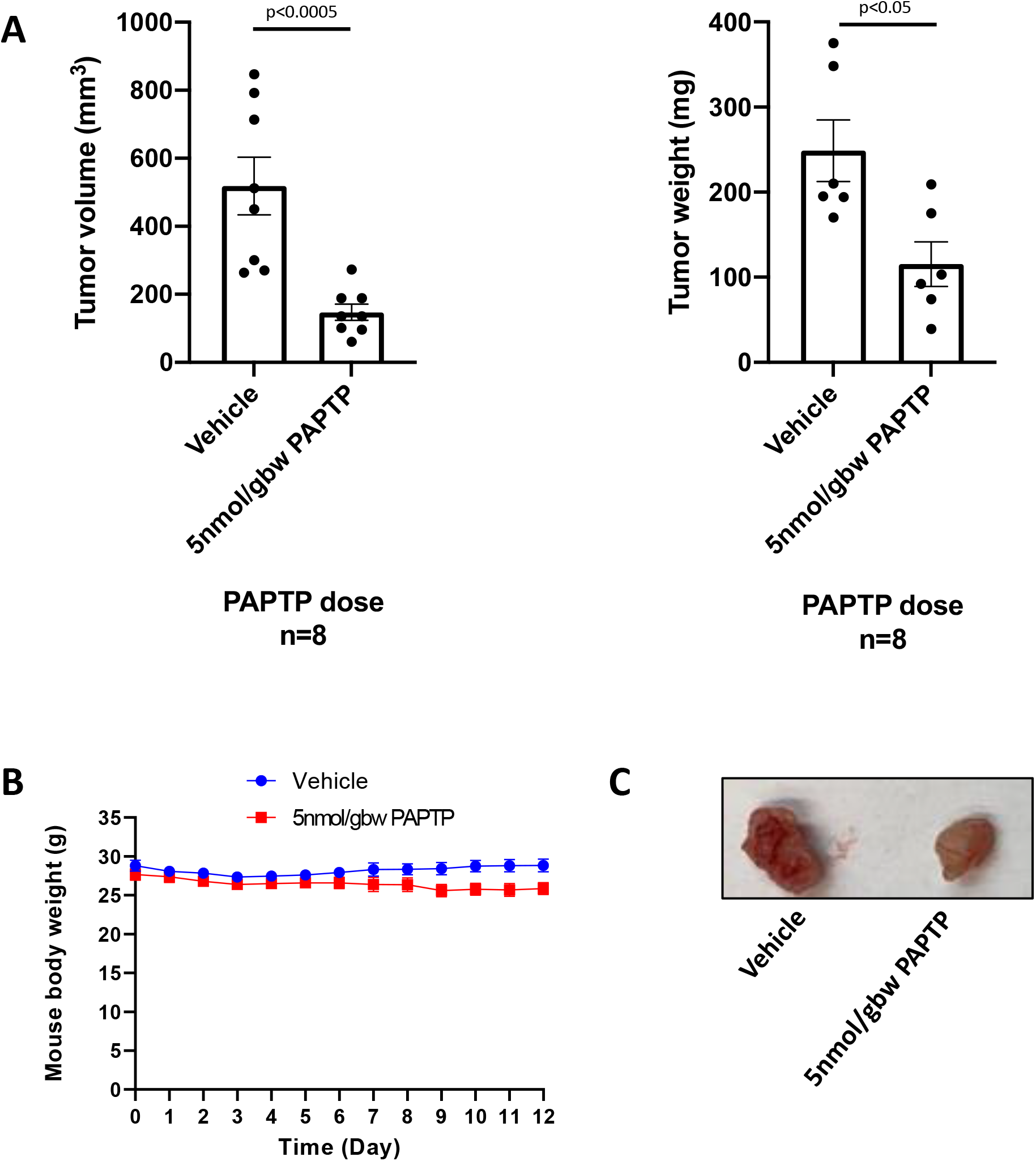
*PAPTP* reduced pancreatic tumor size in mouse models through intraperitoneal administration. A. *PAPTP* reduced mouse pancreatic tumor volume orthotopically significantly by intraperitoneal administration, n=8. *PAPTP* reduced mouse pancreatic tumor weight orthotopically significantly by intraperitoneal administration, n=8. B. Bodyweights change of the mice during *PAPTP* treatment were no difference (p=0.17). C. Representative pictures of the mouse orthotopically injected tumors.

### Concomitant Gemcitabine-Abraxane Treatment with *PCARBTP/PAPTP* Further Reduced Tumor Growth in an Orthotopic Mouse Model

We then examined if there was a synergistic effect of using mitoKv1.3 inhibitors with cytotoxic chemotherapies already used in the clinical practice for pancreas cancer. Compared to untreated controls, the tumor volume and mass reduction seen with 190 nmol/gbw gemcitabine plus 23.4 nmol/gbw Abraxane (albumin-bound paclitaxel) were only 64% and 60% (n=8). However, the tumor volume and mass reduction with 190 nmol/gbw gemcitabine with 23.4 nmol/gbw abraxane plus 15 nmol/gbw *PCARBTP* were 95% and 92%, respectively (n=8). Similarly, we found substantial tumor volume and mass reduction after treatment with 190 nmol/gbw gemcitabine with 23.4 nmol/gbw Abraxane and 5 nmol/gbw *PAPTP* of 80% and 75% (Figure 5).

**Figure 5.**
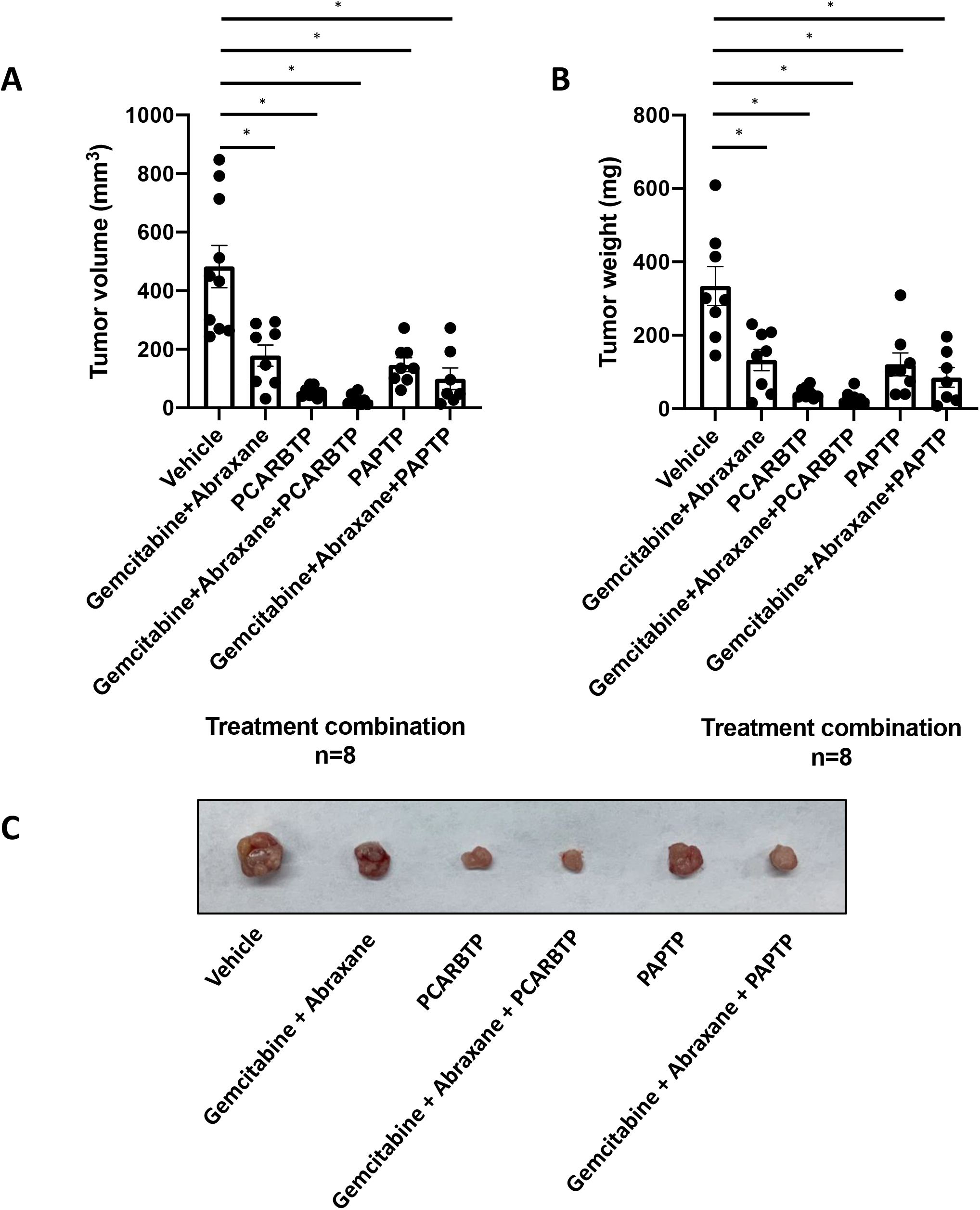
*PCARBTP* or *PAPTP* combined with Gemcitabine and Abraxane reduced pancreatic tumor size and weight in mouse model. A. *PCARBTP* worked better than *PAPTP*, and *PCARBTP* combined with Gemcitabine and Abraxane reduced 95% the pancreas tumor volume. *p<0.0001, compared to the control group. n=8. B. Weight of the tumors same with panel A. *p<0.0001, compared to the control group. n=8. Table 4. Treatment Regimen of Figure 3. C. Representative pictures of the mouse orthotopically injected tumors.

**Table 4.**
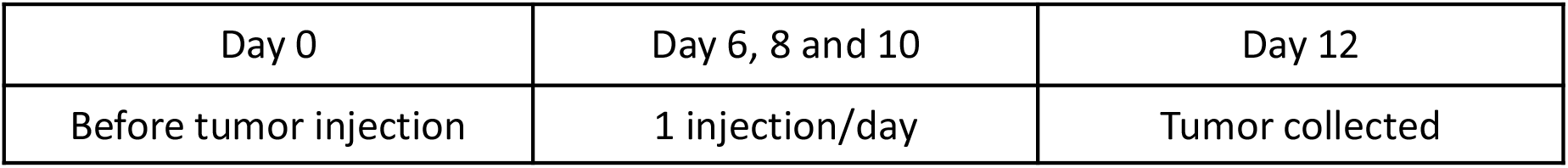
Treatment Regimen of Figure 3. Treatment Regimen of Figure 3.

**Table 5.**
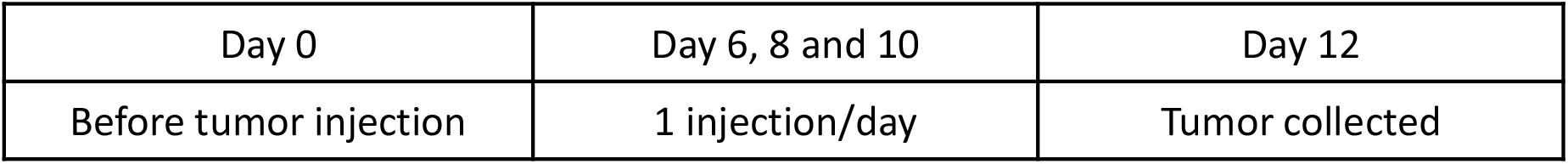
Treatment Regimen of Figure 5B.

### Combined Treatment with Mitochondrial Kv1.3 Inhibitors and Cytotoxic Chemotherapies had No End Organ Toxicities

We have previously reported that *PAPTP* at 5 nmol/gbw and *PCARBTP* at 10 nmol/gbw were not toxic and did not cause changes in the electrocardiogram of mice ^16^. Here we observed that mice did not show sign of distress and show that H&E staining at the time of sacrifice after treatment with the above described combination of *PCARBTP/PAPTP* with Gemcitabine/Abraxane (see concentrations above) showed there was no significant toxicity to the heart, lung, liver, and kidney (Figure 6).

**Figure 6.**
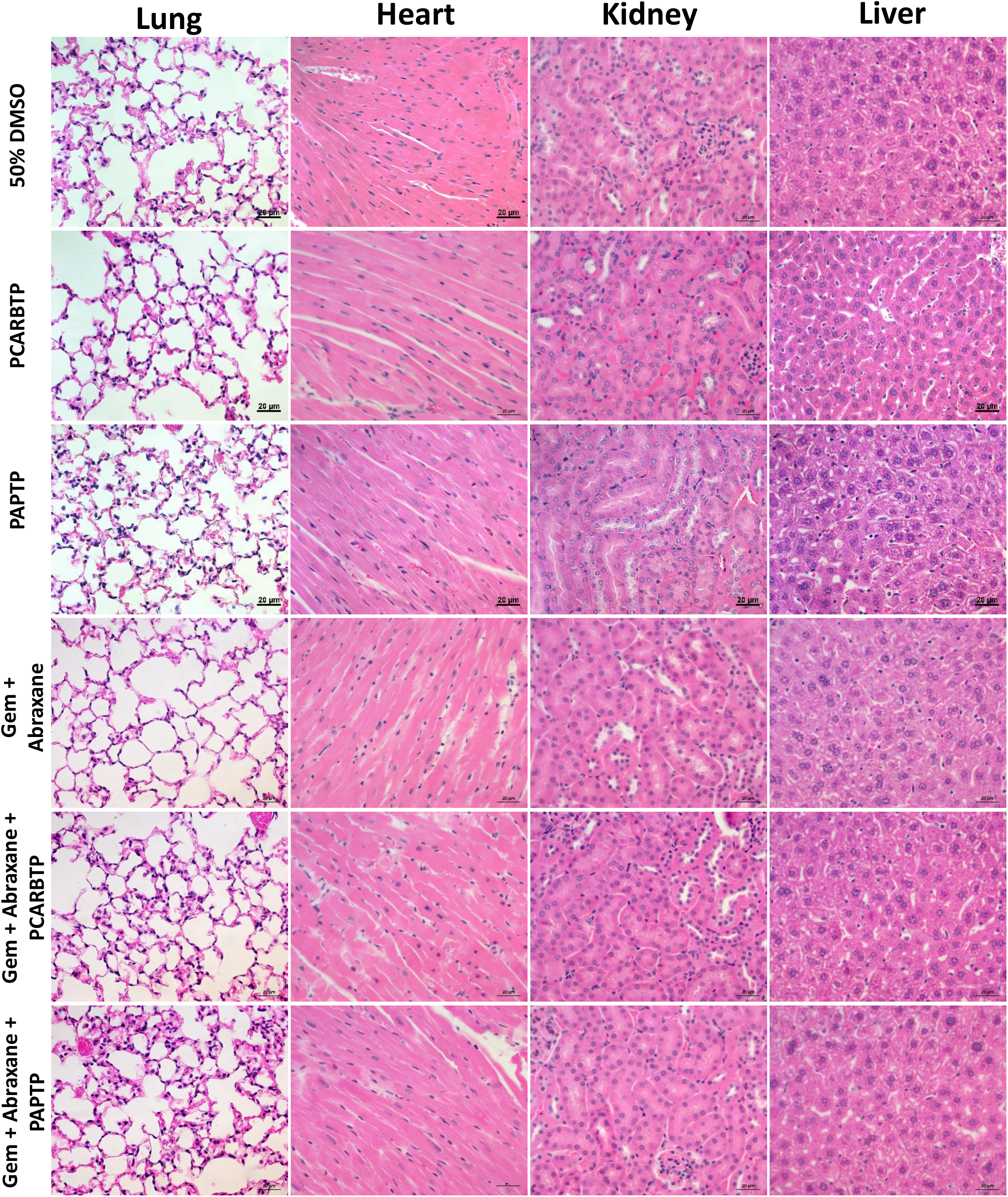
*PCARBTP*/*PAPTP* toxic on other organs in mouse model. H&E staining showed either *PCARBTP* or *PAPTP* intraperitoneal administration only or combined with Gemcitabine and Abraxane had no toxic effective on heart, lung, kidney and liver. Scale bar =20μm.

### Mitochondrial Kv1.3 Inhibitor *PCARBTP* Treatment Activated the Phosphorylation of p38 MAPK, SAPK/JNK

We have previously shown that the action of sub-lethal concentrations of *PCARBTP* was exacerbated when the stress response kinase JNK was inhibited, at least in Jurkat lymphocytes 19. Here, we evaluated the effect of various concentrations of *PCARBTP* on JNK activation monitored by phosphorylation of the kinase, as well as activation of p38 mitogen-activated protein kinase (MAPK), given that both kinases regulate apoptosis induced by several forms of cellular insults. Pan02 cells were treated by *PCARBTP* (up to 10 μM). Western blot analysis with P-p38 MAPK, P-SAPK/JNK showed there was an increase in phosphorylated levels of these proteins when treating the cells with high concentration of the drugs (Figure 7A).

**Figure 7.**
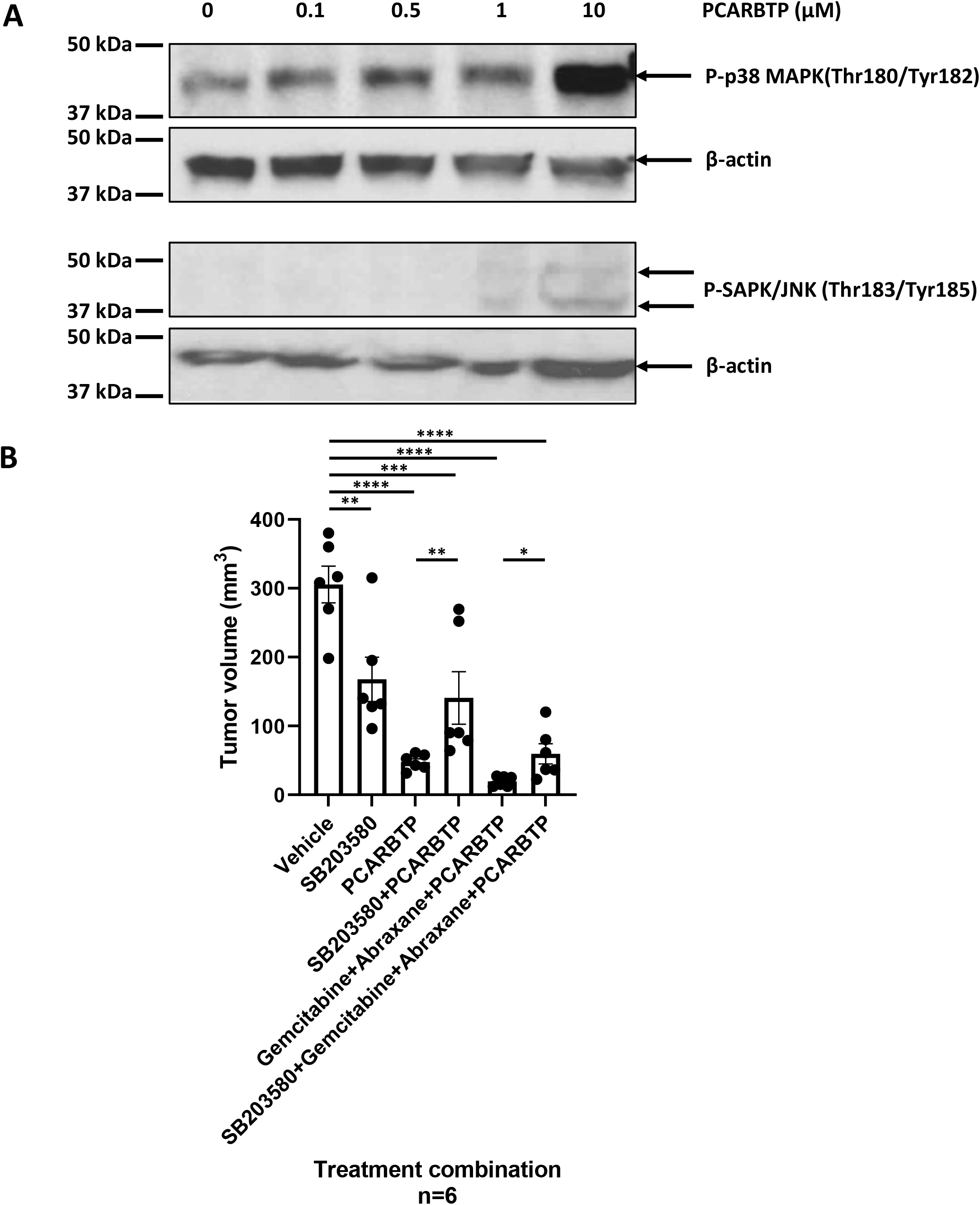
*PCARBTP* mediated the cell apoptosis through the p38 MAPK pathway. A. Phosphorylation levels of p38 MAPK (Thr180/Tyr182), P-SAPK/JNK (Thr183/Tyr185) were increased during the *PCARBTP* treatment for 30 min and by dose-dependent way, n=3. B. p38 MAPK inhibitor (SB203580) attenuated the function of *PCARBTP* and *PCARBTP* combined with Gemcitabine and Abraxane, n=6. *p<0.05; **p<0.01, ***p<0.0005, ****p<0.0001.

To test the significance of p38 MAPK for mitoKv1.3 inhibitor mediated cell death, we examined the effects of treatment with the p38 MAPK inhibitor, SB203580. Treatment with 13 nmol/gbw SB203580 was found to attenuate the effects of *PCARBTP* in an orthotopic mouse model (Figure 7B). Compared to the vehicle control, tumor volume from all drug treated groups were significantly decreased (p<0.05). SB203580 treatment alone reduced tumor volume by 42%. Compared to *PCARBTP* only, the addition of SB203580 to *PCARBTP* decreased growth inhibition by mitoKv1.3 inhibitor treatment (tumor volume 87% vs 51%, p<0.01). Similarly, the addition of SB203580 to Gemcitabine-Abraxane-*PCARBTP* attenuated the tumor-reducing effect of the drugs, as tumor volume was reduced only by 80% (versus 95%) in the presence of p38 MAPK inhibitor (p<0.05), suggesting that *PCARBTP* mediated cell death occurs at least in part through the p38 MAPK pathway, even *in vivo*.

### Resistance to *PCARBTP* Therapy

To examine if resistance to the most efficient mitoKv1.3 inhibitor treatment occurs, we generated a *PCARBTP* resistant clone using the mouse PDAC cell line we used for the *in vivo* studies (Pan02), and compared protein expression to normal Pan02 cells by SWATH nanoLC-MS/MS. The analysis showed that 50 proteins were upregulated and 8 proteins were downregulated. The upregulated proteins were Fasn, Aldh2, Phgdh, Anxa3, Ldha, Aldoa, Ugdh, PRDX6, Hsph1, Pgk1, Tpi1, Gpi1, Krt19, Esd, Gsto1, Psmd2, Aldh3a1, Acly, Ezr, Arf5, Ephx1, Por, G6pdx, LEG3, FRIL1, Serpinb6a, Krt7, Xdh, Vat1, Naca, Gsr, Ywhag, Tfrc, Mif, Pgm1, Akr1c13, Aacs, Gsta2, Plin3, Psph, Hnrnpa1, Ddx39, Pygb, Hars, Eif5, Ranbp3, Hic2, Slc2a1, Uso1, Gng7 (Figure 8A and Table I). As indicated by STRING and pathway enrichment analysis, these proteins grouped into a few main functional classes, linked to metabolism of xenobiotics by cytochrome P450 (Figure 8B), the anti-oxidant defense system, and metabolic pathways of carbohydrates, amino acids and carboxylic acid (Figure 6B). The 8 downregulated proteins identified by proteomics were Anxa2, Anxa1, Vcl, Cand1, Cyb5r3, Dpy30, Slc7a5, Dnajc4 that were not linked to any enriched pathway.

**Figure 8.**
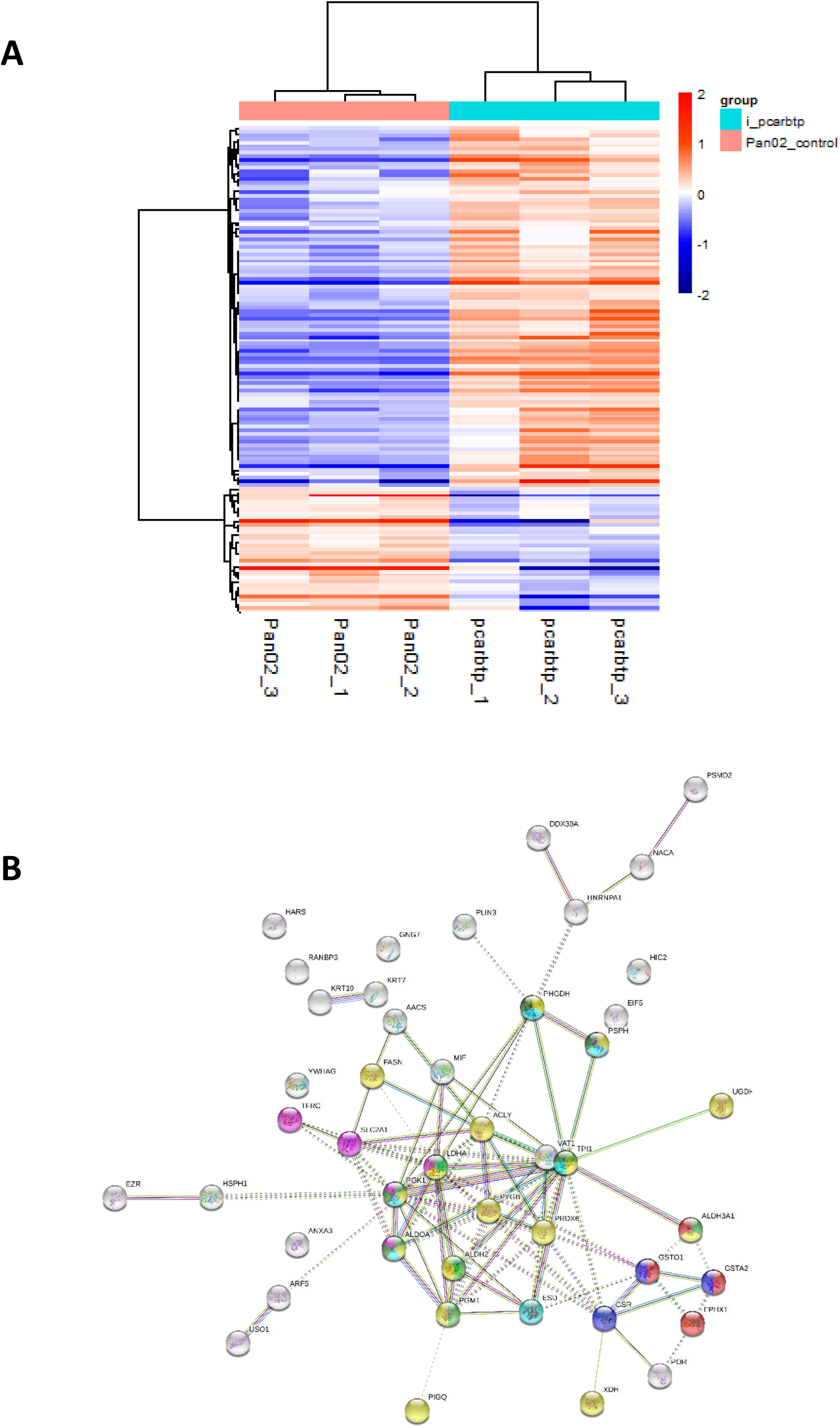
Drug resistance to *PCARBTP* developed by pancreatic tumors A. Proteomic analysis of protein changes in *PCARBTP* resistant clones. Heatmap of unbiased clustering of only the significant proteins, 135 proteins with ANOVA p < 0.05 (color in Log2-scale), n=3. B. STRING analysis showing the following enriched KEGG pathway-linked proteins. Number of nodes: 45; number of edges: 89; average node degree: 3.96; avg. local clustering coefficient: 0.521; expected number of edges: 23; PPI enrichment p-value:< 1.0e-16. KEGG Pathways: Light green: Glycolysis / Gluconeogenesis (<u>hsa00010</u>) (p value 3.45e-08); Yellow: Metabolic pathways (<u>hsa01100</u>) (p value 4.26e-07); Light blue: Carbon metabolism (<u>hsa01200</u>) (p value 1.08e-05); Dark green Biosynthesis of amino acids (<u>hsa01230</u>) (p value 2.10e-05); Magenta: HIF-1 signaling pathway (hsa04066) (p value 7.12e-05); Red: Metabolism of xenobiotics by cytochrome P450 (<u>hsa00980</u>) (p value 0.00039); Dark blue: Glutathione metabolism (<u>hsa00480</u>) (p value 0.0029).

## Discussion

Treatment with mitochondrial Kv1.3 inhibitors *PAPTP* and *PCARBTP* in an orthotopic mouse model using mouse Pan02 cells resulted in tumor growth inhibition of almost 90%. When treatment was combined with cytotoxic chemotherapy Gemcitabine and Abraxane (albumin-bound paclitaxel), an even greater tumor growth reduction of 95% was seen. Most importantly, this treatment strategy did not result in organ toxicity. We are not aware of a similarly successful treatment in PDAC. These studies open the possibility to identify further drug combinations that allow complete eradication of the tumor. We chose to use Gemcitabine and Abraxane as it is one of the common cytotoxic chemotherapy regimens used in the treatment of PDAC and is a 2 drug regimen in comparison to FOLFIRINOX (5-fluorouracil, oxaliplatin, irinotecan), which involves 3 drugs and associated with different toxicity profile.^20, 21^ Although there was further reduction in tumor growth with addition of Gemcitabine/Abraxane, this study showed that there are subpopulations of tumor cells that escape cell death. Most probably, these remaining cells have an upregulated anti-oxidant defense system, as we showed at least in vitro that PCARBTP-resistant cells are characterized by this event.

The role of mitochondrial Kv1.3 in cell proliferation and apoptosis in patients with pancreas ductal adenocarcinoma is not well known. We found that Kv1.3 was highly expressed in resectable huan pancreatic cancer specimens. This is consistent with previous data from us showing by Affymetrix analysis as well as Western blots that Kv1.3 is highly expressed in various, largely chemoresistant human PDAC cell lines harboring p53 mutations (PANC-1, AsPC-1, BxPC-3, Capan-1, Colo-357, MiaPaCa2) ^16^. The channel is present also in the mitochondrial fraction of these cells ^22^.

Our study demonstrates increased expression of Kv1.3 in pancreas cancer. A previous investigation by Bielanska *et al*. showed that Kv1.3 was under-expressed in human PDAC specimens, but these immunohistochemical studies were performed in a very limited sample (n=2) of patients^23^. Another study examined 18 patients and found a correlation between Kv1.3 expression decrease and metastasis ^24^. Please note that both studies were performed on tumor samples from patients with metastatic PDAC, which is biologically different than those with resectable, non-metastatic disease. Given that our study found that Kv1.3 is highly expressed in resected PDAC, it is possible that during the transition to a metastatic phenotype, pancreas cancer cells are able to down regulate Kv1.3, possibly due to methylation of the promoter region of the Kcna3 gene encoding for Kv1.3^24^. The findings reported here are in agreement with previous studies showing that K^+^ channels can promote proliferation^25^, while downregulation of Kv1.3 renders cells resistant to apoptotic stimuli ^26^. In addition, overexpression of the channel in our patient samples suggests Kv1.3 is a novel therapeutic target to treat pancreas cancer. All tumor specimen showed Kv1.3 expression, at least a low general level. Even in those patient samples with low general Kv1.3 expression, there are some tumor cells that express Kv1.3 and are likely to respond to Kv1.3 inhibitors. The nature of these cells remains to be determined. It will be interesting to test whether these specific cells have some characteristics of stem cells. When examining if Kv1.3 expression holds prognostic value, we found no association with overall or recurrence free survival. It is possible that in this patient population with resectable PDAC, compared to metastatic PDAC, alterations at the DNA and/or protein level have not occurred to Kv1.3 to make it a prognostic biomarker. In this study with Western blot analysis, we did not examine mitochondrial Kv1.3 expression but rather that of Kv1.3 also found in all cellular compartments (nuclear membrane, Golgi, plasma membrane and mitochondria). What prognostic value mitochondrial Kv1.3 holds in humans remains unclear.

We also examined if there was an association of Kv1.3 expression with presence of nodal metastases, lymphovascular invasion (LVI), and perineural invasion (PNI) and found there was none (p>0.05). This is likely due to the high rates of PNI (91%), LVI (56%), and positive lymph nodes (84%) in these patients even with localized disease. The predictive value of Kv1.3 expression to *PAPTP* and *PCARBTP* treatment in mice was not tested in this study but we have previously shown a positive correlation between Kv1.3 expression and PCARBTP/PAPTP-induced death in human PDAC lines as well as between expression of the channel in mitochondria and in the plasma membrane.^16^

Mechanistically, the pathway of *PCARBTP* or *PAPTP* induced tumor cell death was associated with ROS production able to drive the cells over a critical point, as indicated by the finding that N-acetyl-cysteine, a molecule able to boost the antioxidant system prevented the *in vivo* tumor reducing effect of both drugs^16^. Here we show that additional mechanisms also come into play, namely an increase in the phosphorylation of p38/MAPK and SAPK/JNK was found. When treated with the p38/SAPK inhibitor, SB203580, there was a reduction in *PCARBTP*-mediated *in vivo* effect, suggesting that mitoKv1.3 inhibition triggers, at least in part, the p38 and/or SAPK-mediated death pathway. These findings are consistent with existing data that show activation of the p38/MAPK and SAPK/JNK pathways to be a favorable prognostic marker and associated with improved overall survival in patients with PDAC.^27–29^

Despite an unprecedented 95% tumor growth inhibition using a combination of mitoKv1.3 inhibitors and cytotoxic chemotherapies, there were still viable tumor cells. After developing a *PCARBTP* treatment resistant cell line, proteomic analysis revealed there was an almost a four-fold increase in the antioxidant system (see Figure 6B). Compared to normal human cells, cancer cells have increased reactive oxygen species (ROS) generation due to reduced ability to produce scavengers.^30, 31^ Inhibition of mitoKv1.3 can initiate a cascade of events that leads to transient hyperpolarization of the inner mitochondrial membrane, formation of ROS, stimulation of permeability transition pore, release of cytochrome c, and resulting apoptosis.^26, 31^. In addition, our recent findings indicate that PAP-1 derivatives bound to mitoKv1.3 are positioned in a way that their psoralenic moieties can directly accept electrons from complex I of the respiratory chain and donate these electrons to molecular oxygen, further boosting ROS.^18^ Given the importance of ROS generation to the mechanism of mitoKv1.3 inhibitor mediated cell death, it is not surprising to see that cancer cells resistant to *PCARBTP* upregulated the antioxidant system (GSR, GSTO1, GSTA2, for description of function see Table I). Likewise, it is not surprising that proteins linked to the cytochrome P450-mediated detoxification system are upregulated (EPHX1, GSTO1, GSTA2, ALDH3A1), as this system is mainly responsible for xenobiotic metabolism in the cells ^32^. In addition, an upregulation of proteins linked to metabolic processes such as glycolysis, carboxylic acid biosynthesis and amino acid synthesis can be observed (see Figure 6D). This result can be interpreted as a consequence of the inhibition of the mitochondrial respiration and ATP production ^16^. In this respect it is interesting to note that the metabolic effects of the Shk-186 toxin, that acts only on the plasma membrane located Kv1.3^33^ are different from those reported here for inhibition of mitoKv1.3.

Future studies should focus on using mitoKv1.3 inhibitors to selectively kill cells with higher Kv1.3 expression, which may result in not only the direct killing of cancer cells but also enhance the activity of cytotoxic chemotherapies and potentially immunotherapies.

## Conclusion

The potassium channel Kv1.3 is found to be overexpressed in pancreas adenocarcinoma and mitoKv1.3 inhibitors combined with cytotoxic chemotherapies can result in greater than 95% tumor growth reduction without organ toxicity. *In vitro* western blot analysis and *in vivo* orthotopic model results indicated *PCARBTP*-mediated cell death indeed occurs through the p38 MAPK pathway. SWATH-MS and STRING and pathway enrichment analysis found the proteins linked to anti-oxidant defense system, and metabolic pathways of carbohydrates, amino acids and carboxylic acid are different in *PCARBTP* resistant clones. These data suggest utilizing Kv1.3 should be considered a novel therapeutic target for pancreatic cancer treatment.

## Materials and Methods

### Orthotopic Mouse Pancreatic Tumor Injection Model

All animal experiments were approved by University of Cincinnati Ethic Committee and Institutional Animal Care and Use Committee. Eight-week-old, wild type male, C57BL/6J mice were purchased from Jackson Labs (000664, Jackson Labs, USA). Mice were anesthetized using 120mg/kg ketamine plus 20mg/kg xylazine. Orthotopic injection was performed as described by Tepal *et al.^34^* In detail, a left subcostal incision was made just below rib cage and the pancreas was identified. The tumor cell suspension was created by mixing 25μL of Matrigel with 25μL of Pan02 cells (National Cancer Institute- Frederick Cancer Research and Development Center, Frederick, MD, USA) containing 1×10^6^ cells. Pan02 cells were cultured in DMEM+10% FBS medium, under 37℃ and 5% CO_2,_ no antibiotic added. The tumor suspension was slowly injected into the pancreas and the needle left in place for 60 seconds to allow the Matrigel to set. After ensuring hemostasis, the abdomen was closed in 2 layers using 3-0 silk suture.

### *In Vivo* Kv1.3 Inhibitor and Cytotoxic Chemotherapy Administration

*PCARBTP* was employed with two kinds of application methods. For Figure 2 A, B, C and D, *PCARBTP* was suspended in 50% DMSO at concentration of 25 mM and 50 mM and filled in 1003D mini-pumps (MICRO-OSMOTIC PUMP MODEL 1003D, Reservoir Volume, 100 μl, 1.0 μl per hour, 3 days). The doses of PCARBTP equated to 1nmol/h/gbw and 2nmol/h/gbw. The mini-pumps were implanted into the peritoneal cavity on day 6 after tumor injection. Pumps were replaced after drug release was completed. In the remaining experiments, *PCARBTP* was suspended in 50% DMSO and injected into the peritoneal cavity at a dose of 15 nmol/gbw on day 6 after tumor injection. Similarly, *PAPTP* was administrated at a dose of 5 nmol/gbw. Gemcitabine was dissolved in ddH_2_O and injected into the intraperitoneal cavity at dose of 190 nmol/gbw 6 days after tumor injection. Abraxane was dissolved in DMSO and intraperitoneally injected at a dose of 23.4 nmol/gbw.

### Orthotopic Pancreatic Tumor Transplantation Model

Mice were injected pancreatic tumor as described as above, then treated with 15 nmol/gbw *PCARBTP* or 5 nmol/gbw *PAPTP* at day 6, 8, and 10 post tumor injection for 3 doses. Mice were kept until day 21 and the pancreatic tumors were isolated and digested with 0.25% trypsin-EDTA (25200056, Thermo Fisher, USA) and DNase I (DN25, Sigma, USA), single cells were collected, 1×10^6^ primary cells in 25μL were mixed with 25μL of Matrigel and slowly injected into the pancreas of wild type C57BL/6J mice (000664, Jackson Labs, USA) as described above. The 2^nd^ cycle mice were treated with 15nmol/gbw *PCARBTP* or 5nmol/gbw *PAPTP* at day 6, 8, and 10 post tumor injection and kept until day 21 after which tumors were collected for analysis.

### Immunohistochemistry

Immunohistochemistry staining was performed using a biotin-streptavidin-peroxidase (SP) kit (AB64269, Abcam, USA) and a diaminobenzidine kit (DAB) as previously described^19^. Institutional Review Board approval was obtained to obtain human resected pancreas ductal adenocarcinoma specimens and associated clinicopathologic data (IRB 2019-0324). Tumor specimens were obtained from the University of Cincinnati Department of Pathology. Five-micrometer sections were deparaffinized and rehydrated in xylene and gradients of ethanol. Slides were boiled in citrate buffer (10mM sodium citrate, 10mM citric acid, pH 6.0) at 92–98 ℃ for 10 min to retrieve the antigen. The sections were then incubated with 3% H_2_O_2_ in methanol for 10 min to quench endogenous peroxidase and blocked with normal goat serum for 20 min. Sections were incubated with specific primary antibodies against Kv1.3 (1: 200; P4497, Sigma, USA) at 4℃ overnight. The sections were then incubated with biotinylated goat-anti-rabbit IgG secondary antibody and sections stained with DAB working reagent (per manufacturer’s instructions) for 30–60 seconds. They were then counterstained with hematoxylin. Finally, sections were mounted with Permount (SP15-500, Fisher, USA) onto slides. Negative control was performed using unconjugated rabbit IgG (011-000-003, Jackson ImmunoResearch Laboratories, Inc., West Grove, PA, USA). Images were acquired with a ZEISS AXIO microscope (Carl ZEISS, Germany). Slides were scored by a gastrointestinal pathologist who specializes in evaluating pancreas cancer. He was blinded to all clinicopathologic data and determined percent and intensity of staining. A final score of high versus low expression were obtained based on these data.

### Western Blot

Cells were lysed in whole-cell lysis buffer (50 mM HEPES, 150 mM NaCl, 1 mM EGTA, 10 mM sodium pyrophosphatate, 1.5 mM MgCl_2_, 100 mM NaF, 10% glycerol and 1% Triton X-100, pH 7.2) containing an inhibitor cocktail (1mM phenylmethylsulfonyl fluoride, 10 mg/ml aprotinin and 1mM sodium orthovanadate) to extract total protein. Protein concentrations were determined using a standard bicinchoninic acid (BCA) assay (23225, Thermo Fisher Scientific Inc., Waltham, MA, USA), and 50 μg of total protein was subjected to 10% SDS-PAGE followed by electrotransfer onto nitrocellulose membranes. The membranes were blocked in 5% skim milk, and then incubated overnight at 4 ℃ with primary antibodies against human Kv1.3 (1: 1000, P4497, Sigma, USA), P-p38 MAPK (1: 1000; Cell signaling), P-SAPK/JNK (1: 1000; Cell signaling), β-actin (1: 2 000; Abcam). Membranes were then washed with TBST for 3×10 min. This was followed by incubation with horseradish peroxidase-conjugated secondary antibodies for 1 h at room temperature in 5% skim milk and washed with TBST for 3×10 min. Immunoreactive signals were detected using enhanced chemiluminescence (Pierce, Rockford, IL, USA). Three independent experiments were performed.

### H&E Staining

Slides containing paraffin sections were passed using the following steps: 3 × 5min in Xylene (blot excess xylene before going into ethanol), 2 × 5min in 100% ethanol, 1 × 5min in 95% ethanol, 1 × 5min in 70% ethanol, 1 × 5min deionized H_2_O, and 1 × 3min Hematoxylin. They were then rinsed with deionized water 1 × 5min, tap water and ethanol to destain. After subsequent rinse, they were treated 1 × 30 sec Eosin, 3 × 5min 95% ethanol, 3 × 5min 100% ethanol, 3 × 5min Xylene, and coverslip was placed using Permount mounting medium (SP15-500, Fisher, USA).

### *PCARBTP* Resistant Clones

Pan02 cells were detached from tissue culture flasks before reaching confluence by removing the culture medium, adding trypsin-EDTA and incubating for 3 min at 37℃ and 5% CO_2_. After this incubation period, fresh medium was added and cells were spun at 800 rpm for 5 min. Supernatant was removed and fresh medium added. Cell count was carried out with the standard trypan blue exclusion method. Pan02 cells were seeded at 1 cell per well in 400 μL of fresh medium in 96 flat-bottomed well plates. Clones were inspected regularly so those wells with more than 1 clone could be discarded. The addition of *PCARBTP* was carried out by replacing medium containing *PCARBTP* at the different doses. If the cells survived at that dose for more than 3 days, we increased the dose, and obtained the 4 clones of Pan02 cells that survived under 10μM *PCARBTP* medium. Resistant cells were amplified under 10 μM *PCARBTP* then proteins were collected for proteomic analysis.

#### Proteomic Analysis of Resistant Clones

The Pierce 660nm Protein assay was performed on a 1:10 dilution of the samples to determine the protein concentration using BSA as a standard. Sufficient protein was present such that 50 μg was taken out to run on a short 1D gel for digestion. 50 μg of each sample (non-resistant and resistant clones) in 40 μL of Laemmli buffer were run 1.5 cm into a 1D, 1.5 mm 4-12% BT gel using MOPS running buffer. Pre-stained protein markers were used in surrounding lanes. The regions between the markers and the dye front were excised for trypsin digestion following the standard in gel protocol. The resulting peptides were extracted, dried, and prepared for mass spectrometry. 2.5 μg of each sample was run on the nanoLC-MS/MS in DDA mode and the combined DDA runs were searched using Protein Pilot (SCIEX, AB Sciex Pte. Ltd, USA) to create the protein spectral library. 720 proteins were identified with 99% confidence with an FDR of less than 1% at peptide and protein level. A matched SWATH-MS method in DIA mode of the samples was used to collect quantitative data for each of the samples for the comparative profiling. Three clones of *PCARBTP* resistant cells and three replicates of normal Pan02 were processed. SWATH-D data analysis workflow was used to validate the data set and detected significant quantitative changes.

### Statistical Analysis

Clinicopathologic data were obtained from the electronic medical records. Variables include patient age, gender, histologic grade, lymphovascular invasion, perineural invasion, stage, lymph node status, and oncologic status. Statistical analyses were performed using SPSS 26 (IBM Inc., Armonk, NY). Descriptive statistics are reported as the median and interquartile range (IQR). Chi-squared tests or Fisher’s exact test were used for comparing categorical variables. Median values or continuous variables were examined using Student’s t-test or Mann-Whitney. Survival analyses were performed using Kaplan-Meier methodology. P-values <0.05 were considered statistically significant. The tumor volume and mass and mouse body weight were analysed by t-test and one-way ANOVA using GraphPad Prism 9.0, each experiment has more than 6 mice in one group indicated in Results and Figure legends. P<0.05 was considered as statistically significant. *p<0.05; **p<0.01, ***p<0.0005, ****p<0.0001.

## Acknowledgments

The authors would like to thank the Boyce Family Foundation for their support in this research endeavor. This study was funded, in part, by the Central Surgical Association Turcotte Award, 2019 and in part by the Italian Association for Cancer Research (IG 20286). The authors thank Dr.s Lucia Biasutto, Sofia Parrasia and Andrea Rossa for help with chemical analysis of the K^+^ channel inhibitors used in this study and Dr. Mario Zoratti for useful discussion and critical reading of the manuscript.

## Authors’ Contributions

**Conception and design:**WL, MB, MJE, IS, CP, AM, EG, SAA, and SHP

**Development of methodology:**WL, MB, MJE, GCW, CP, AM, JW, IS, EG, SAA, and SHP

**Acquisition of data (provided animals, acquired and managed patients, provided facilities, etc.):**WL, MB, GCW, CP, AM, JW, EG, SAA, and SHP

**Analysis and interpretation of data (e.g., statistical analysis, biostatistics, computational analysis):**WL, MB, MJE, GCW, JW, CP, AM, IS, EG, SAA, and SHP

**Writing, review, and/or revision of the manuscript:**WL, MB, MJE, GCW, CP, AM, JW, IS, EG, SAA, and SHP

**Administrative, technical, or material support (i.e., reporting or organizing data, constructing databases):**WL, EG, and SHP

**Study supervision:**WL, MB, MJE, GCW, EG, SAA, and SHP

**Other (provided intellectual input and edited the article):**WL, MB, MJE, GCW, CP, AM, JW, IS, EG, SAA, and SHP

## The Paper Explained

### Problem

Treatment of pancreas ductal adenocarcinoma (PDAC) remains challenging due to the late stage of presentation, limited efficacy of cytotoxic chemotherapies, and aggressive tumor biology. Novel therapeutic targets are desperately needed. The voltage-gated potassium channel, Kv1.3, is one such unique target. It has been extensively studied in many cancers but less is known in pancreas cancer. In this study we evaluated tissue expression of Kv1.3 in resected PDAC and tumor inhibition using novel Kv1.3 inhibitors developed by our group (*PCARBTP* and *PAPTP*).

### Results

We found that in the largest cohort of surgically resected PDAC specimens, Kv1.3 exhibits high tissue expression in over half of tumors. Utilizing orthotopic pancreas tumor mouse models, treatment with mitochondrial Kv1.3 inhibitors, PCARBTP and PAPTP resulted in tumor reductions of 87% and 70%, respectively. When these inhibitors were combined with cytotoxic chemotherapies (gemcitabine/abraxane), this resulted in a 95% tumor growth inhibition without organ toxicity. Evaluation of the mechanism of cell death using *in vivo* models showed an integral role of the p38-MAPK pathway. Finally, resistant clones to mitochondrial Kv1.3 inhibitors escaped cell death through upregulation of the anti-oxidant system.

### Impact

The findings from this study indicate that Kv1.3 is expressed in early stage, non-metastatic, resectable pancreas cancer specimens. Treatment with novel mitochondrial Kv1.3 inhibitors resulted in 95% reduced tumor growth when combined with cytotoxic chemotherapies. This near complete eradication of tumors using this treatment strategy shows that Kv1.3 represents an innovative therapeutic target for pancreas cancer therapy.

---

**Table I.**
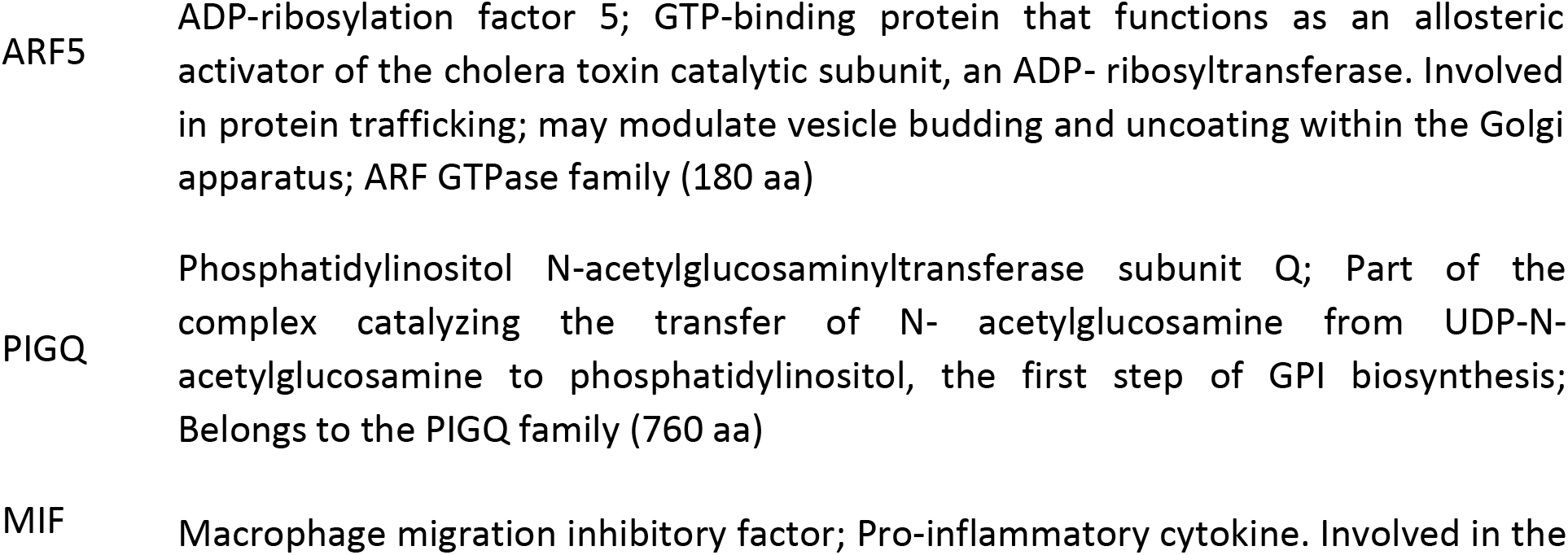
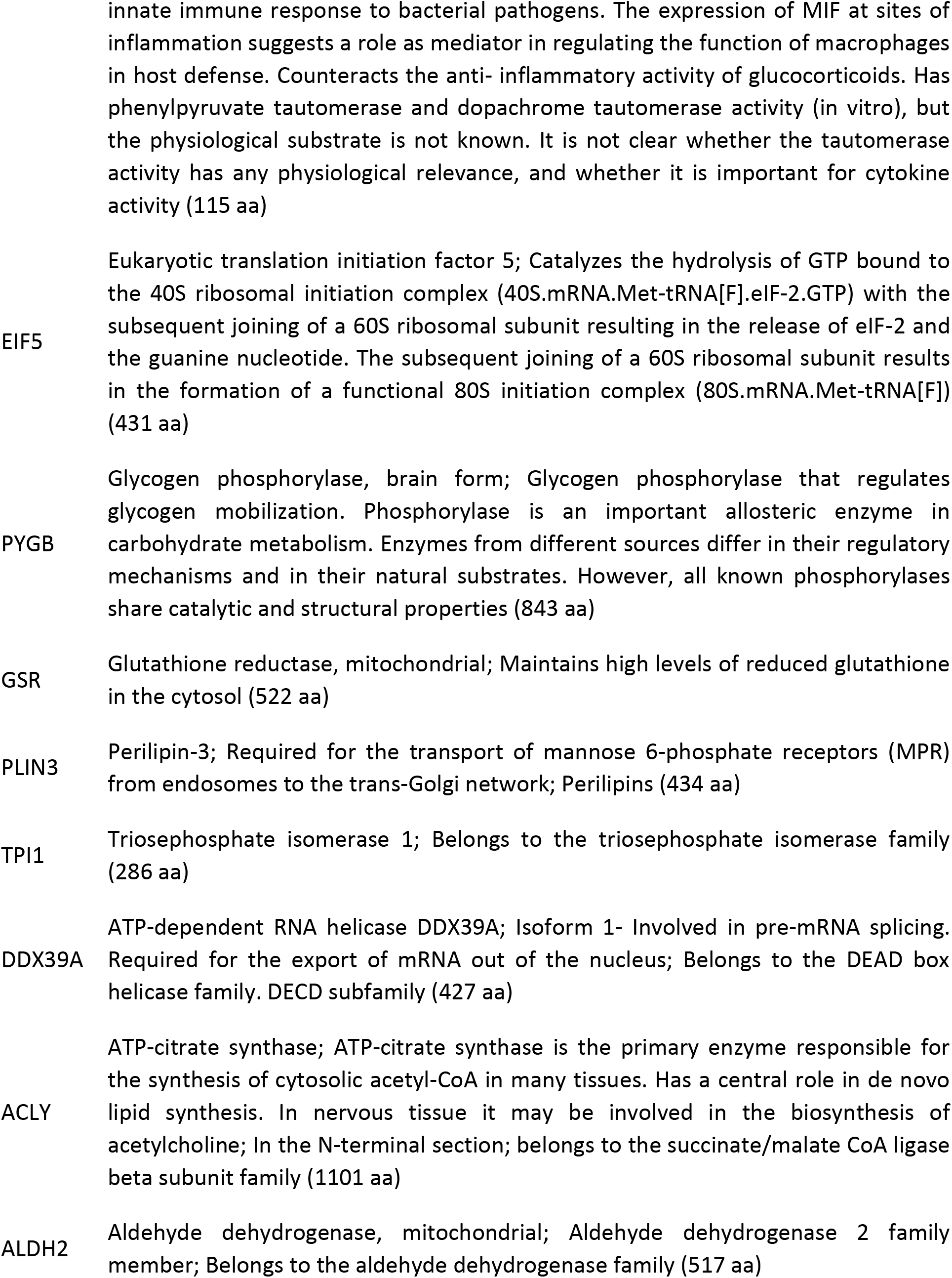
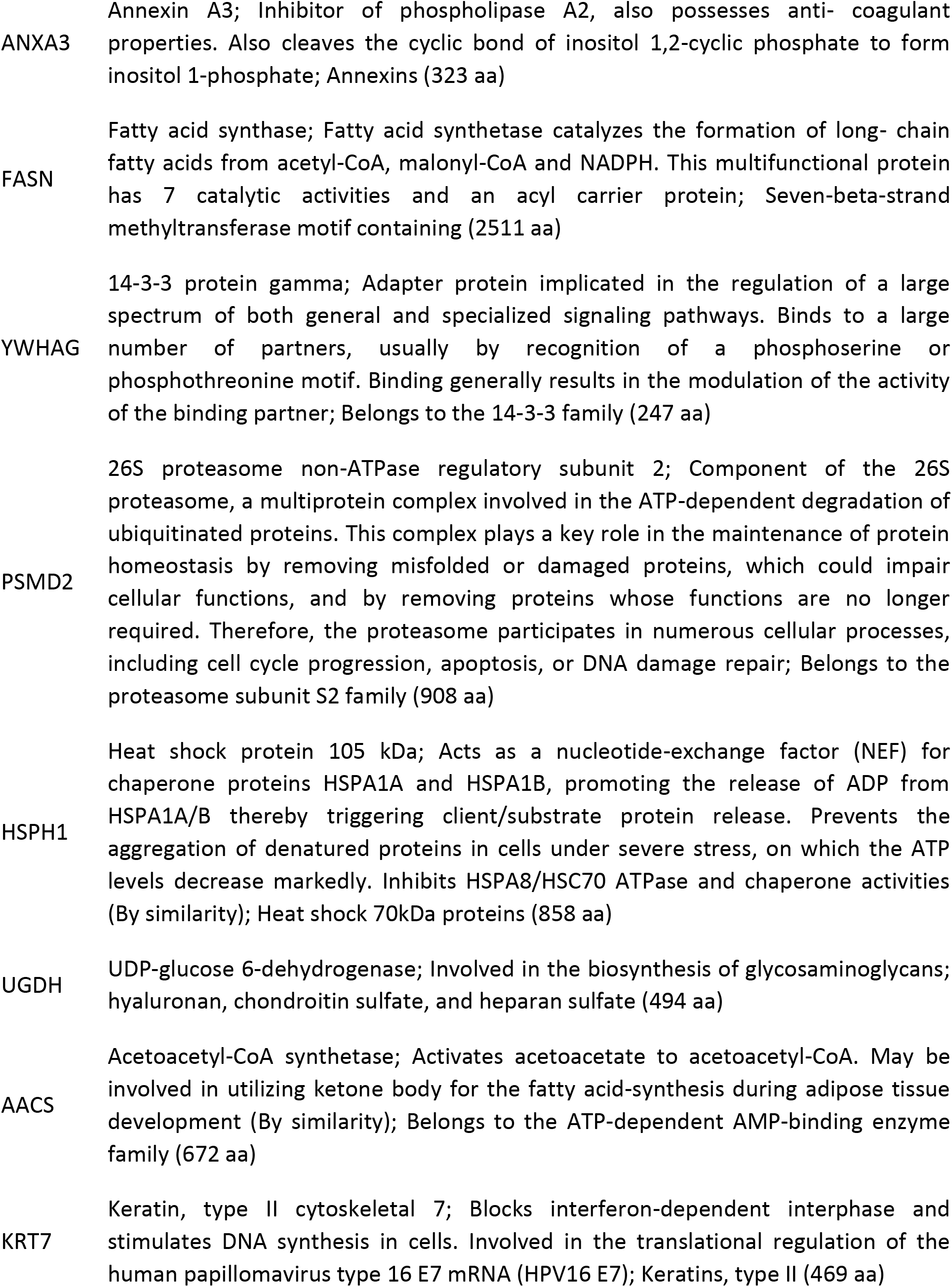
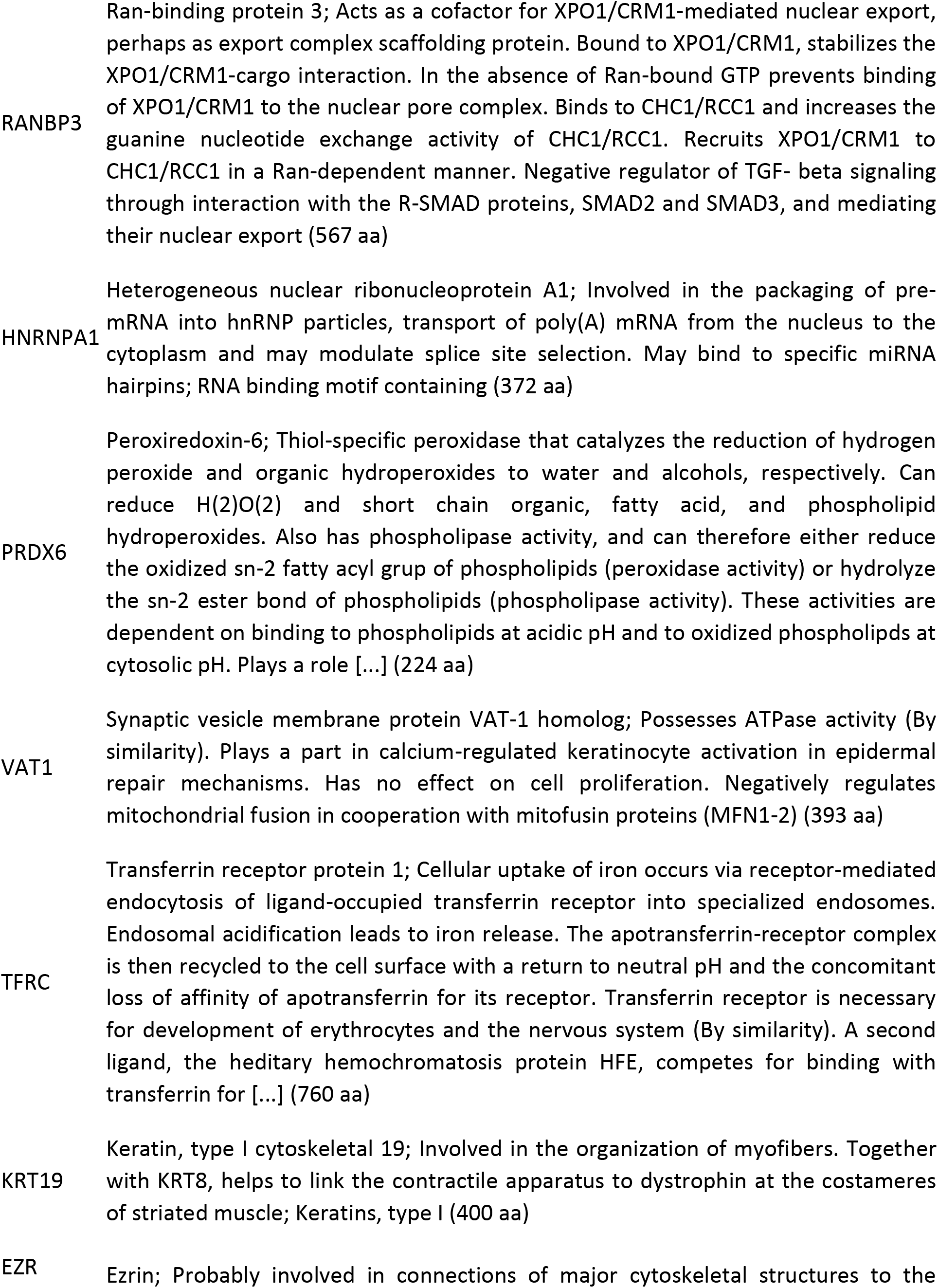
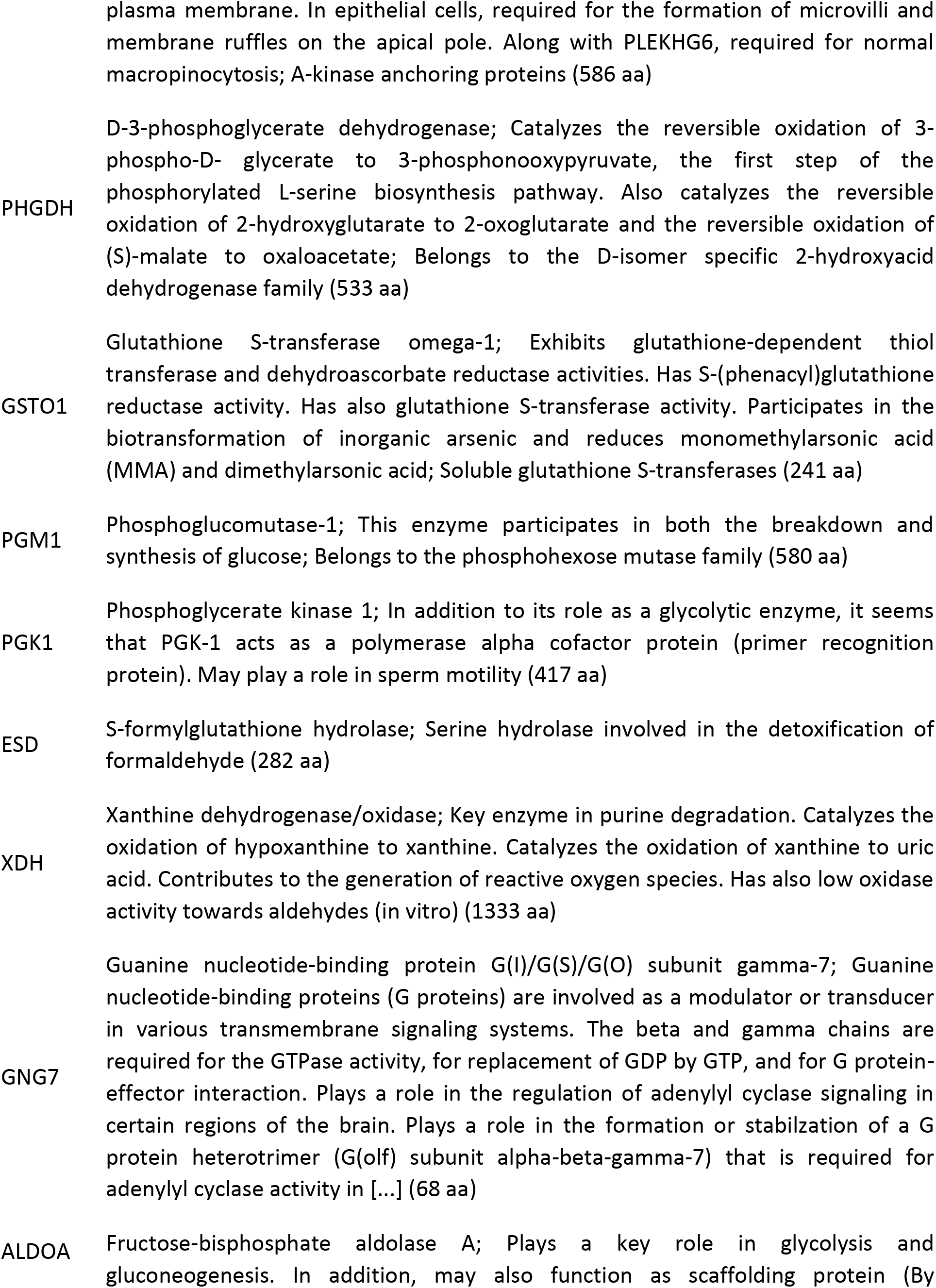
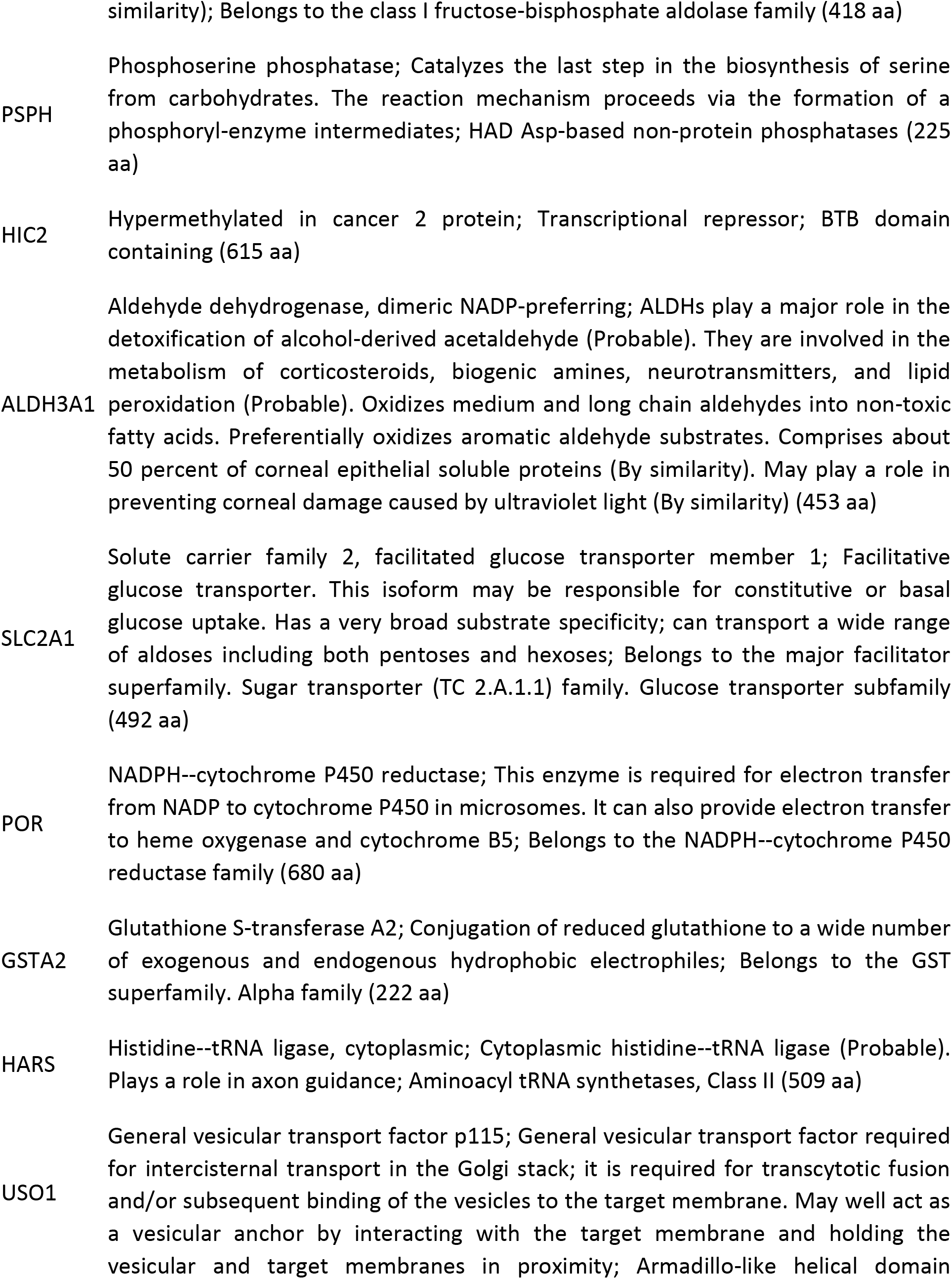
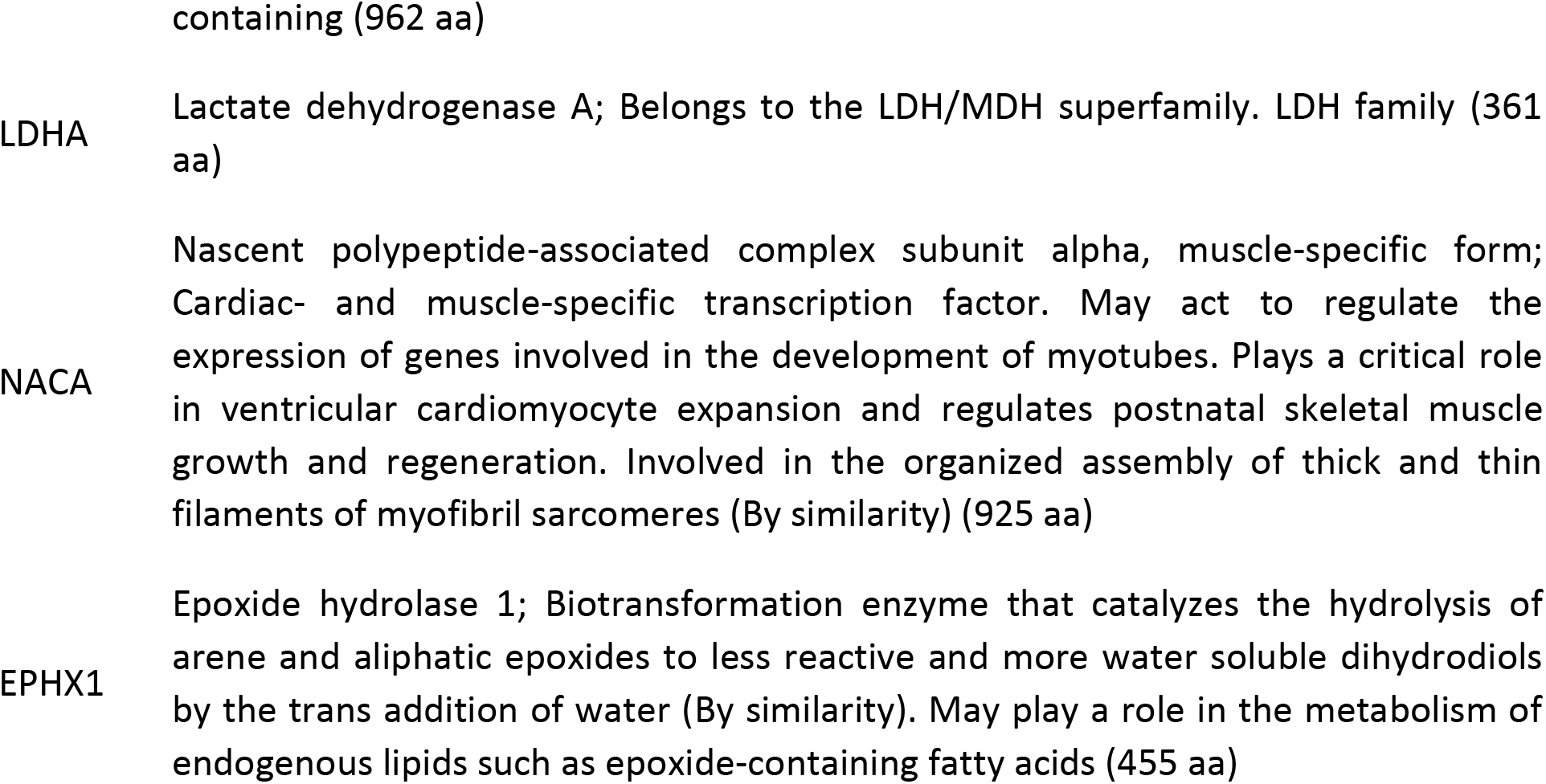
SWATH nanoLC-MS/MS analysis of protein expression in *PCARBTP*-resistant Pan02 clones compared to normal Pan02 cells. List of upregulated proteins and their function. Source: STRING analysis (https://stringdb.org/cgi/input.pl?taskId=_notask&sessionId=SF25ZH1QUDpd)

